# A genetic strategy to allow detection of F-actin by phalloidin staining in diverse fungi

**DOI:** 10.1101/2025.04.29.651289

**Authors:** Alison C.E. Wirshing, Analeigha V. Colarusso, Daniel J. Lew

## Abstract

Actin is highly conserved across eukaryotes. This versatile protein builds cytoskeletal networks central to diverse cellular processes, including cell division and cell motility. The most potent and broadly used reagents to detect polymerized actin distribution in fixed cells are fluorescently conjugated derivatives of the basidiomycete-derived toxin, phalloidin. However, despite its conservation, actin in many ascomycete fungi fails to bind phalloidin. Here we trace the failure to bind phalloidin to a single amino acid change in a phalloidin-binding residue in actin. Reverting this change in the fungus *Aureobasidium pullulans* by introducing the point mutation *act1*^*V75I*^ at the native *ACT1* locus confers phalloidin binding without disrupting actin function. We describe a simple genetic technique to introduce this point mutation that may be effective in other fungal systems. This strategy should enable characterization of F-actin in a wider range of fungi.

**SUMMARY:** A single point mutation, *act1*^*V75I*^, enables phalloidin staining in ascomycete fungi

## INTRODUCTION

Actin networks drive diverse fundamental cellular processes including cell motility and morphogenesis, vesicle traffic, endocytosis, organelle positioning, and cytokinesis (1). The molecular mechanisms by which actin cytoskeletal networks are assembled have been elucidated, in part, through studies in the tractable yeast model systems *Saccharomyces cerevisiae* and *Schizosaccharomyces pombe* (2–4). This work has established several core functions of the actin cytoskeleton in fungi, but these simple small yeasts do not engage in the more elaborate forms of fungal morphogenesis like building mycelial networks and fruiting bodies. Thus, studies of broader fungal morphological and biological diversity exploit other species like *Aspergillus nidulans*, whose long hyphae provided a model for investigating mechanisms of long-range intracellular transport (5), and *Magnaporthe sp*., whose appressorium served as a model for the highly pressurized structures used by fungi to penetrate plant hosts (6). New genome modification technologies have accelerated the process of making nonmodel organisms tractable for molecular and cellular biology, broadening the range of biological behaviors that can be probed at the mechanistic level (7, 8). One example of this expansion is the polymorphic fungus, *Aureobasidium pullulans*, which is emerging as a model for how the same cytoskeletal networks are repurposed to grows cell of remarkably different shapes and sizes (9–12).

One of the challenges to investigating the roles of actin is that for some fungi, phalloidin staining is ineffective (6, 13–15). Phalloidin is a phallotoxin produced by the death cap mushroom *Amanita phalloides*, that binds with high specificity to F-actin allowing detection of F-actin networks in diverse cell types with high contrast and low background signal (16–18). While there are alternative approaches to visualizing actin, these do not provide comparable sensitivity or spatial resolution. In addition, other methods may compromise aspects of actin function. Tagging actin directly with a fluorescent tag compromises its function, and use of fluorescently tagged actin binding proteins can result in poor decoration, and therefore detection, of subsets of actin networks (19). Thus, phalloidin remains the gold standard for visualizing cellular actin networks, and the failure of some fungal actins to bind phalloidin has hindered progress in those systems. Here we find that *A. pullulans* actin does not bind phalloidin, and that a single nucleotide change in actin (I75V) is shared among fungi whose actin fails to bind phalloidin. By introducing a V75I point mutation at the native actin gene locus, we show that this single mutation is sufficient to allow phalloidin to bind *A. pullulans* F-actin, without detectably compromising actin function. A similar approach would likely be successful in other fungal species.

## RESULTS

### Fungal species whose actin fails to bind phalloidin have a single key reside in common

Phalloidin staining fails to label F-actin in several ascomycete fungi including *Aspergillus nidulans, Magnaporthe oryzae, Neurospora crassa*, and *Trichoderma citrinoviride* (6, 13–15). These fungi all belong to the largest ascomycete subdivision, pezizomycotina. However, F-actin is successfully detected by phalloidin in ascomycete species in saccharomycotina (e.g., *Geotrichum candidum, Saccharomyces cerevisiae*, and *Ashbya gossypii*) and more distant taphrinomycotina (e.g., *Schizosaccharomyces pombe*) subdivisions (13, 20–22). This suggests that changes to actin resulting in failure to bind phalloidin may have occurred relatively recently in fungal evolution, perhaps being selected for toxin resistance in species that share a habitat with *A. phalloides*. To identify relevant differences in actin between the species whose actin can and cannot bind phalloidin, we aligned each actin sequence and focused on residues shown to mediate phalloidin binding though structural and genetic studies (**Figure 1**) (23, 24). We included *Gallus gallus* (chicken) actin because the crystal structure of *G. gallus* actin bound to phalloidin was recently published (24). This structure showed that residues E72, H73, I75, T77, L110, N111, P112, R177, D179 of the first actin monomer (closer to the barbed end) and T194, G197, Y198, S199, F200, E205, and L242 of the second actin monomer in a filament interact with phalloidin (24). Many of the key residues that mediate phalloidin binding (e.g., I75, T77, L110, and T194) vary among fungi, but only residue 75 correlates with the ability to bind phalloidin. Residue 75 encodes isoleucine in species where actin binds phalloidin, and valine in species where it does not (**Figure 1**). We therefore reasoned that this single amino acid change could underly loss of actin binding to phalloidin.

**Figure 1:**
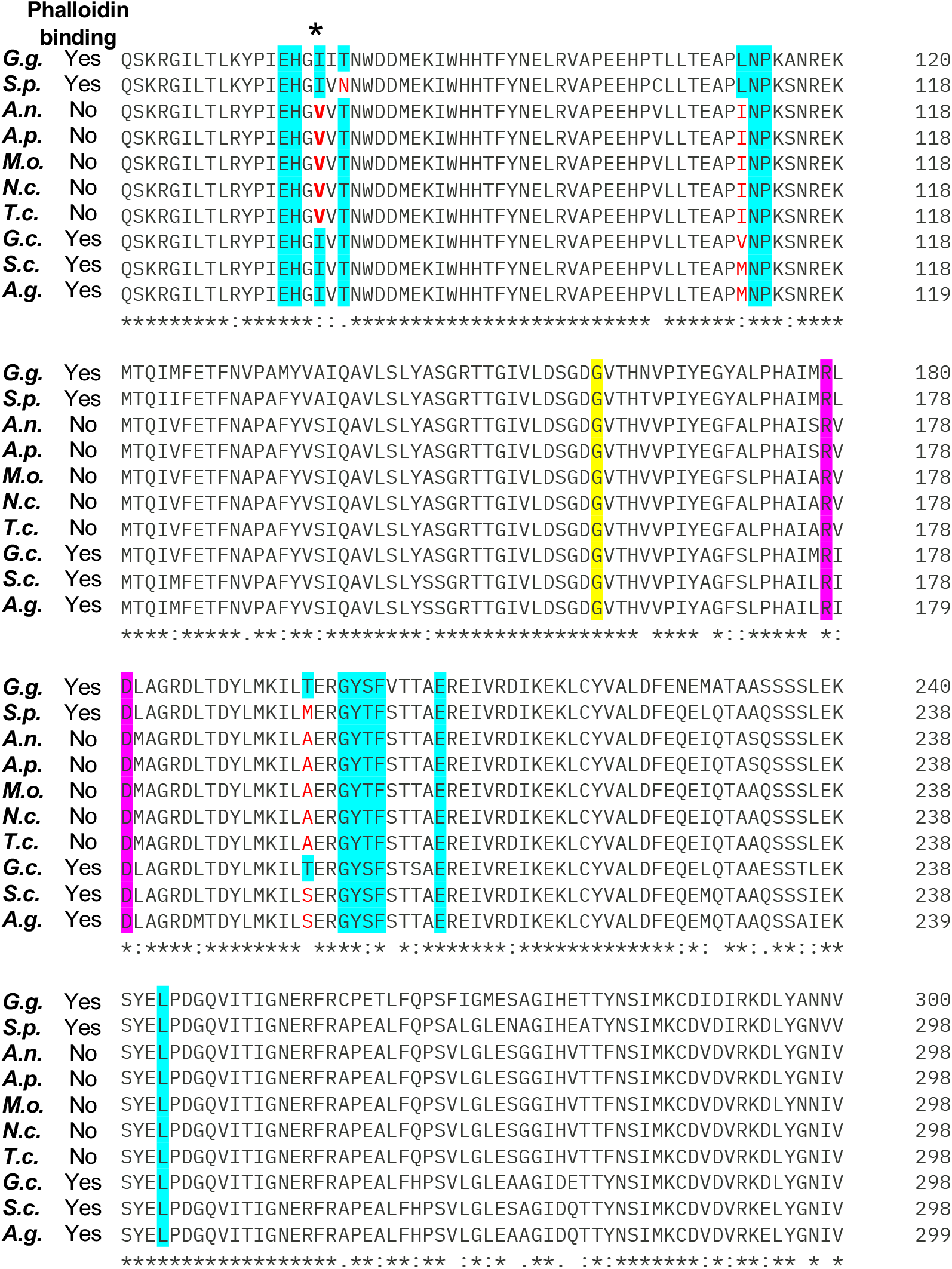
Alignment of the actin protein sequences from species whose actin does (“Yes”) or does not (“No”) stain with phalloidin. The GenBank accession number for each sequence is in parenthesis: *Gallus gallus* (G.g., yes, NP_001385203.1), *Schizosaccharomyces pombe* (S.p., yes, NP_595618.1), *Aspergillus nidulans* (A.n., no, XP_664146.2), *Aureobasidium pullulans* (A.p., no, XP_029758358.1), *Magnaporthe oryzae* (M.o., no, XP_003719871.1), *Neurospora crassa* (N.c., no, XP_011394625.1), *Trichoderma citrinoviride* (T.c., no, XP_024751298.1), *Geotrichum candidum* (G.c., yes, KAF5095009.1), *Saccharomyces cerevisiae* (S.c., yes, NP_116614.1), *Ashbya gossypii* (A.g., yes, NP_983171.1). Numbers in the figure indicate sequence position. Residues important for phalloidin binding based on structural data are highlighted in cyan, based on genetics in *S. cerevisiae* are highlighted in yellow, and from both are highlighted in magenta (23, 24). Cases where these important residues are not conserved are in red. The only key residue that is different in all of the non-phalloidin-binding actin sequences is indicated with an asterisk. Sequence alignment generated using Clustal Omega (41).

### Scarless replacement of *ACT1* with *act1*^*V75I*^

Like other fungi in pezizomycotina, *A. pullulans* has a valine at position 75 in actin, and our preliminary attempts to visualize F-actin with phalloidin staining in *A. pullulans* were unsuccessful (see **Figure 3E**). To introduce a V75I point mutation at the native *ACT1* locus, we built a plasmid to replace native *ACT1* with *act1*^*V75I*^ using a ‘pop-in/pop-out’ strategy. This strategy takes advantage of efficient homologous recombination to first insert *act1*^*V75I*^ next to endogenous *ACT1*, and then remove the wild type *ACT1*, yielding a precise scarless gene replacement (**Figure 2**). Insertion and removal are selected for using the counter-selectable marker *UR*A3 (25). *URA3* enables synthesis of uracil, but it also converts the non-toxic compound 5-fluoroorotic acid (5-FOA) to the toxic metabolite 5-fluorouracil. Thus, cells with a wildtype *URA3* gene grow on media lacking uracil but die on media containing 5-FOA, while cells that lack *URA3* cannot grow on media lacking uracil but do grow on media containing 5-FOA.

**Figure 2:**
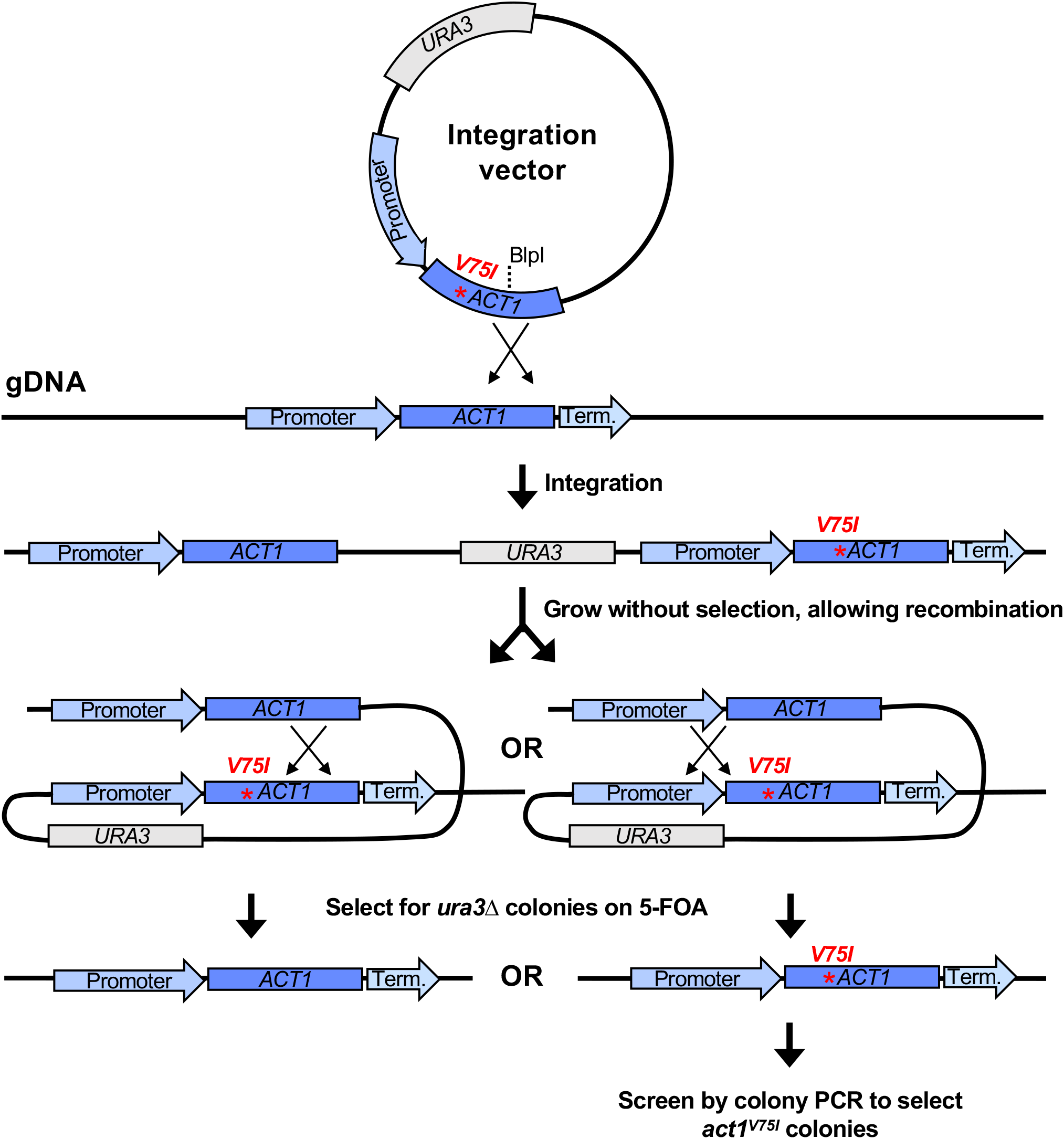
Schematic showing the ‘pop-in/pop-out’ gene replacement strategy. Linearization of the integration vector targets integration at the native *ACT1* locus. This results in the chromosomal *ACT1* and *act1*^*V75I*^ sequences flanking *URA3* and the vector backbone. Colonies that have undergone homologous recombination to ‘pop-out’ *URA3* are selected for on 5-FOA. Recombination downstream of the V75I point mutation pops out *act1*^*V75I*^ while recombination upstream of the V75I point mutation pops out *ACT1* leaving a scarless *act1*^*V75I*^ mutation. The *act1*^*V75I*^ pop-outs are then identified by colony PCR.

**Figure 3:**
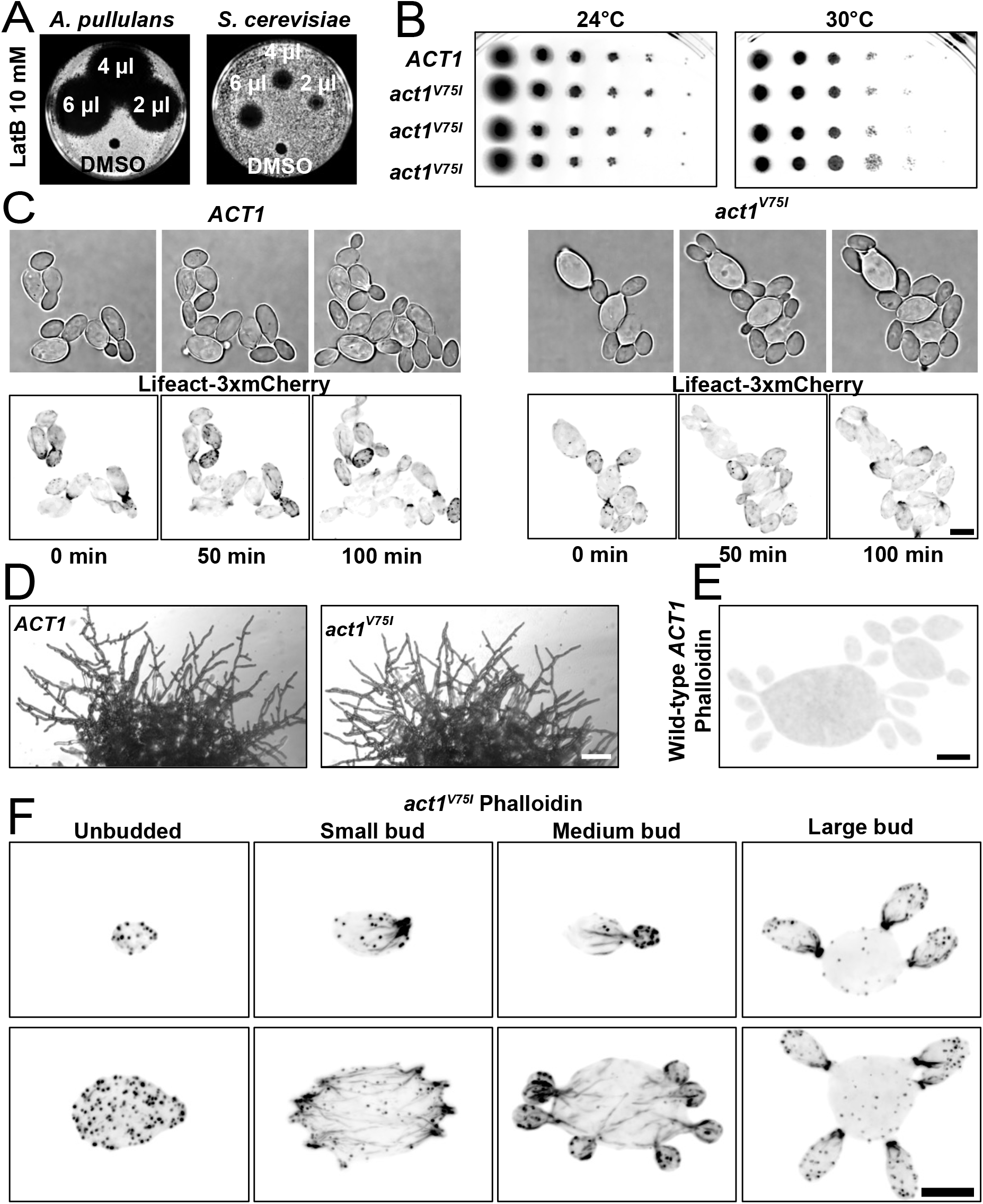
A single point mutation in actin is sufficient to allow phalloidin binding without altering cell growth. (A) Lawns of wild-type *A. pullulans* (DLY23540) or *S. cerevisiae* (DLY22370) cells spotted with the indicated amount of LatB (10 mM) or the carrier control (DMSO). Dark areas indicate lack of cell growth. (B) Growth of 10-fold serial dilutions of wild-type cultures (DLY23540) or cultures with the *act1*^V75I^ point mutation (DLY24813, DLY24814, and DLY24816) after 2 days of growth on YPD at the indicated temperatures. (C) Brightfield and maximum intensity projections of images of live wild-type (DLY24628) and *act1*^V75I^ (DLY24872) cells expressing the F-actin marker Lifeact-3xmCherry. Scale bar 5 µm. (D) Images of 2 day old colonies of *ACT1* (DLY23540) and *act1*^V75I^ (DLY24813) cells. Scale bar 100 μm. (E) Maximum intensity projection of a wild-type, *ACT1*, cell (DLY23540) fixed and stained with phalloidin showing no detection of F-actin networks. (F) Maximum intensity projection images of *act1*^V75I^ cells (DLY24813) of different sizes and bud numbers fixed and stained with phalloidin. Phalloidin decorates small dots, patches, and linear cables in small and medium budded cells. Large-budded cells show enrichment of actin at the neck. Scale bar 5 µm.

We previously reported a *ura3ΔHYG*^*R*^ strain of *A. pullulans* (9). Into this strain, we transformed our plasmid with the *A. pullulans URA3* and *act1*^*V75I*^ genes (**Figure 2**). Digestion at the unique BlpI site was used to target integration of the plasmid to the *ACT1* locus (**Figure 2**), and transformants were selected on media lacking uracil. Following integration, transformants have two copies of the actin gene, *ACT1* and *act1*^*V75I*^, separated by the *URA3* gene and plasmid backbone (**Figure 2**). Growth on rich media allows rare recombination events to ‘popout’ *URA3* (**Figure 2**). Pop-outs were recovered by selection on media containing 5-FOA. When recombination occurs upstream of the V75I point mutation, a scarless *act1*^*V75I*^ replacement is introduced at the native locus (**Figure 2**). Successful *act1*^*V75I*^ colonies were identified by colony PCR followed by sequencing.

### *act1*^*V75I*^ allows for phalloidin staining of F-actin without compromising fungal growth or morphology

As with other fungi, F-actin is necessary for growth of *A. pullulans*, and depolymerization of F-actin with latrunculin B (LatB) blocks growth (**Figure 3A**). In contrast, *act1*^*V75I*^ mutant cells grow similarly to wild-type even at an elevated temperature, showing that the *act1*^*V75I*^ allele functions to support cell growth (**Figure 3B**).

*A. pullulans* exhibit yeast, hyphal, and meristematic growth morphologies (26, 27). In their yeast form, cells can be multinucleate and generate multiple buds in a single cell cycle (9, 11, 12). *act1*^*V75I*^ mutants were morphologically similar to wild-type, capable of growing multiple buds in yeast form (**Figure 3C and F**) as well as hyphae at colony borders (**Figure 3D**). Although much weaker than phalloidin staining, actin networks can be visualized using fluorescent Lifeact (an F-actin binding peptide) in *A. pullulans* (11). In an attempt to improve actin labeling, we tagged Lifeact with three tandem copies of mCherry codon-optimized for *A. pullulans*. This strategy works very well in *S. cerevisiae* (28) but was less effective in *A. pullulans* (**Figure 3C**), highlighting the need to develop species-specific tools for F-actin detection. Regardless, Lifeact-decorated F-actin appeared similar in *act1*^*V75I*^ mutants and wildtype cells (**Figure 3C**). Together, these findings indicate that the V75I substitution does not appreciably affect actin function in *A. pullulans*.

We next tested if *act1*^*V75I*^ improved the ability to detect F-actin in *A. pullulans* using phalloidin staining. Formaldehyde fixation and staining of wild-type cells results in no visible F-actin staining (**Figure 3E**). However, when the same staining protocol was applied to *act1*^*V75I*^ cells, phalloidin decorated F-actin networks similar in appearance to the dense Arp2/3-generated endocytic patches and linear formin-assembled actin cables seen in other fungi (**Figure 3F**) (3). In large-budded cells, F-actin accumulated at the mother-bud neck, potentially forming an actomyosin ring for cytokinesis (**Figure 3F**) (29). Thus, introducing a single point mutation in *ACT1* is sufficient to enable phalloidin staining to detect F-actin in *A. pullulans*.

## DISCUSSION

We find that a phalloidin interacting residue in actin, I75, is changed to V75 in ascomycete fungi whose actin fails to bind phalloidin. Introducing a V75I point mutation is sufficient to enable phalloidin staining of F-actin in *A. pullulans* without compromising actin function. This strategy to enable phalloidin staining will likely be applicable to other ascomycete fungi. For example, *A. nidulans* and *A. fumigatus* have identical actin sequences to *A. pullulans*. Thus, the same point mutation is expected to enable phalloidin staining of F-actin in these fungi. Further, it has been shown that *S. cerevisiae ACT1* supports growth of *A. fumigatus* (14). As our approach requires only a single amino acid change, it limits the likelihood that this mutation will interfere with actin function in other systems. Our genetic strategy relies on 5-FOA counterselection of *URA3+* cells, which has been shown to work in a wide range of fungi (25, 30–35). Precise gene replacement also relies on efficient homologous recombination, which is also found in many fungi. For species that preferentially use nonhomologous end joining, mutation of end joining factors has proven effective in increasing the frequency of homologous recombination (36–38). Thus, with minimal modifications we hope that this strategy will enable phalloidin detection of F-actin in a wider range of fungi, expanding the tools available for investigating the roles of the actin cytoskeleton in diverse biological contexts.

## METHODS

### *A. pullulans* strains and maintenance

All experiments were conducted with *Aureobasidium pullulans* strain EXF-150 and derivatives (39). The exception is the LatB sensitivity assay where wild-type *S. cerevisiae*, YEF473 background (DLY22370), was also used (40). *A. pullulans* strains used in this study are as follows: wild-type (DLY23540), *ura3ΔHYG*^*R*^ (DLY24148), *ura3ΔHYG*^*R*^; *act1*^*V75I*^ (DLY24813, DLY24814 and DLY24816), *URA3:Lifeact-3xmCherry* (DLY24628), and *URA3:Lifeact-3xmCherry*; *act1*^*V75I*^ (DLY24872). Unless otherwise indicated, *A. pullulans* was grown at 24°C in standard YPD medium (2% glucose, 2% peptone, 1% yeast extract) with 2% BD Bacto™ agar (214050, VWR) in plates.

### Generation of *A. pullulans act1*^*V75I*^ mutants

The *act1*^*V75I*^ point mutation was introduced using plasmid DLB4812. This plasmid has 1212 bp upstream of the *ACT1* (protein ID 347440) start codon and the mutant *act1*^*V75I*^ coding sequence. The point mutation was introduced using primers. These sequences were introduced into the plasmid backbone pAP-U2-1 (Addgene # 236472) (10). This backbone includes the native *URA3* gene under the native URA3 promoter with the *scTEF1* terminator. Linearization with BlpI cuts in the *act1*^*V75I*^ coding sequence, enabling integration (‘pop-in’) at the native *ACT1* locus.

An additional selection step was used to ‘pop-out’ the *URA3* cassette to generate the scarless *act1*^*V75I*^ mutation. After transformation with linearized DLB4812 and selection of uracil prototrophs, transformants were grown overnight at 24°C in 5 ml YPD. These transformants have wild-type *ACT1* and the mutant *act1*^*v75i*^ flanking the *URA3* cassette. To select for uracil auxotrophs that flipped out the *URA3* cassette, 10^6^ cells were plated on 5FOA plates (6.71 g/L BDDifcoTM Yeast Nitrogen Base without Amino Acids, BD291940, FisherScientific, 0.77 g/L Complete Supplement Mixture minus uracil, 1004-100, Sunrise Science Products, 2% glucose, 50 mg/L uracil, 1 g/L 5-Fluoroorotic acid, F10501-25.0, Research Products International, and 2% BD Bacto™ agar, 214050, VWR) and grown for 3 days at 24°C. Colonies growing on 5-FOA were checked by colony PCR to confirm introduction of the V75I point mutation using the Phire Plant Direct PCR Master Mix (F160S, Fisher Scientific) and the amplified *ACT1* (or *act1*^*V75I*^) coding sequence was checked by sequencing (Quintara Bio).

### Generation of *A. pullulans act1*^*V75I*^ mutants with Lifeact-3xmCherry

To visualize F-actin with Lifeact, we used plasmid DLB4810. This plasmid contains the native *URA3* coding sequence and Lifeact-3xmCherry driven by the *scACT1* promoter. This plasmid was built by using primers to introduce the Lifeact sequence into plasmid backbone pAP-U2-3 (Addgene ID# 236474) after removal of the *apACT1* promoter and 3xGFP sequences (10). Plasmid DLB4810 was designed to rescue uracil auxotrophy and integrate at the native *URA3* locus in an auxotrophic background, *ura3ΔHYG*^*R*^ (9). Restriction digest with ApaI and NruI results in a linear cassette flanked by homology to the native *URA3* locus. Transformants restore uracil prototrophy and insert Lifeact-3xmCherry at *URA3*.

All PCR fragments were amplified with Phusion™ Hot Start Flex 2X Master Mix (M0536L, NEB) following the manufacturer’s instructions, and the plasmids were built using NEBuilder HiFi DNA Assembly Master Mix (E2621L, New England Biolabs). Plasmid were confirmed by whole plasmid sequencing (Plasmidsaurus, Eugene, OR, USA).

### *A. pullulans* transformation

*A. pullulans* was transformed using PEG/LiAc/ssDNA as described previously (9). Briefly, cells were grown to a density of ∼10^7^ cells/ml in YPD (4% glucose), harvested by centrifugation, and rinsed with sterile water followed by competence buffer (10 mM Tris pH 8, 100 mM LiOAc, 1 mM EDTA, 1 M sorbitol). Cells were resuspended in competence buffer in a final concentration of 2×10^9^ cells/ml in a final volume of 50 µl (∼10^8^ cells in each tube). At this stage, competent cells were stored in the -80°C freezer or used directly by adding 2.5 µl 40% glucose, 10 µl carrier DNA (10 mg/ml single-stranded fish-sperm DNA, 11467140001, Roche), 15 µl of transforming DNA (linearized plasmid ∼1,000 ng/µl), and 600 µl of transformation buffer (10 mM Tris pH 8, 1 mM EDTA, 40% (w/v) PEG 3350). Cells were incubated in transformation buffer for 1 hour at 24°C while rotating. Following the 1-hour incubation, 30 µl 40% glucose was added, and the cells were heat shocked for 15 min at 37°C. Cells were centrifuged and the transformation buffer was removed. For selection of prototrophs on media lacking uracil, cells were resuspended in 250 µl 1 M sorbitol and spread onto two drop out uracil plates, (6.71 g/L BDDifcoTM Yeast Nitrogen Base without Amino Acids, BD291940, FisherScientific, 0.77 g/L Complete Supplement Mixture minus uracil, 1004-100, Sunrise Science Products, 2% glucose, and 2% BD Bacto™ agar, 214050, VWR).

### Live-cell imaging

For imaging experiments, a single colony was used to inoculate 5 ml of YPD (2% glucose). Cultures were grown overnight at 24°C to a density of 1-5×10^6^ cells/ml. Cells were pelleted at 9391 rcf for 10 s and resuspended at a final density of ∼7×10^7^ cells/ml. Approximately 2×10^5^ cells were mounted on a 8-well glass-bottomed chamber (80827, Ibidi) and covered with a 200 µl 5% agarose (97062-250, VWR) pad made with CSM (6.71 g/L BD Difco™ Yeast Nitrogen Base without Amino Acids, BD291940, FisherScientific, 0.79 g/L Complete Supplement Mixture, 1001-010, Sunrise Science Products, and 2% glucose). All experiments were conducted at room temperature (20-22°C).

Growth of wildtype and *act1*^*V75I*^ cells expressing Lifeact-3xmCherry was monitored by imaging on a Nikon Ti2E inverted microscope with a CSU-W1 spinning-disk head (Yokogawa), CFI60 Plan Apochromat Lambda D 60x Oil Immersion Objective (NA 1.42; Nikon Instruments), and a Hamamatsu ORCA Quest qCMOS camera controlled by NIS-Elements software (Nikon Instruments). Z-stacks (17 slices, 0.7 µm interval) were acquired using 50 ms exposure at 15% laser power (excitation 561 nm).

### Imaging colony morphology

Do determine if the *ac1*^*V75I*^ mutation alters colony morphology, *ACT1* and *ac1*^*V75I*^ colonies were used to inoculate 5 ml YPD and grown overnight at 24°C. The following day, 100-200 cells were plated on CSM plates supplemented with 10% (v/v) YPD and 7% agarose. Plates were grown for 2 days at 24°C. Colony edges were imaged at room temperature on a MSM 400 tetrad dissection microscope (Singer Instruments) equipped with PixeLINK PL-D752MU-T CMOS camera using a 4x objective (NA. 0.10, Singer Instruments).

### Imaging phalloidin-stained F-actin

Wild-type or *act1*^*V75I*^ cells were grown in YPD as in live imaging experiments. Cells were fixed by adding formaldehyde (CAS # 50-00-0, ThermoFisher) at a final concentration of 3.7% to 500 µl cells in YPD. Cells were fixed for 40 minutes at 24°C with agitation. Following fixation, cells were rinsed three times with PBS and resuspended in 30 µl PBS with 0.1% Trition X-100. The Alexa Flour™ 488 phalloidin (A12379, ThermoFisher) stock was prepared in anhydrous dimethyl sulfoxide (DMSO, 66 µM final concentration) and 1.5 µl was added to the 30 µl cells. Cells were incubated with the phalloidin at 24°C with agitation for 2 hours in the dark, rinsed once with 100 µl PBS, mounted in SlowFade™ Glass Soft-Set Antifade Mountant (S36917, ThermoFisher), and imaged immediately. We found that more rinses with PBS or allowing the samples to sit at room temperature for more than one hour prior to imaging decreased the image quality. Fixed and stained cells were imaged on the same spinning disk confocal system used for live cell imaging described above. The entire cell volume was acquired using 75 Z-slices (at 0.2 μm step intervals). Exposure times of 50 ms at 50% laser power (excitation 488 nm) were used.

### Image processing

Confocal images were denoised using DenoiseAI and maximum intensity projections were generated in NIS Elements (Nikon Instruments).

### Growth assay

To compare cell growth of *A. pullulans* strains, a single colony was inoculated into 5 ml of YPD (2% glucose) and grown at 24°C for 48 h. Cultures were serially diluted five times in sterile deionized water in a 96-well plate and transferred onto solid YPD plates using a pin-frogger. Plates were grown for 48 h at 24°C and imaged on an Amersham Imager 680 (General Electric Company) using the colorimetric Epi-white settings.

To determine if depolymerization of F-actin disrupts growth of *A. pullulans*, wild-type *A. pullulans* and *S. cerevisiae* cells were inoculated and grown in YPD as described above. 10^5^ cells from the overnight cultures were spread on YPD plates and briefly allowed to dry. The indicated amount of Latrunculin B stock (10 mM in DMSO) was spotted onto the plates along with a spot of DMSO as negative control. Plates were grown for 2 days at 24°C and imaged as above.

## ABBREVIATIONS

5-FOA: 5-Fluoroorotic acid
DNA: Deoxyribonucleic acid
F-actin: Filamentous actin
Hyg: hygromycin B

## DATA AVAILABILITY

The data generated in this study are available from the corresponding author upon reasonable request.

## ACKNOWLEDGEMENTS

We would like to thank Dr. Miguel A. Peñalva for helpful discussions. This work was funded by NIH/NIGMS grant R35GM122488 to DJL.

## AUTHOR CONTRIBUTIONS

Conceptualization, review, and editing manuscript— A.C.E. Wirshing, and D.J. Lew. Data curation, investigation, methodology, visualization, validation, and formal analysis—A.C.E. Wirshing and A. Colarusso. Drafting of manuscript— A.C.E. Wirshing. Project administration, supervision, and resources— D.J. Lew.

